# Establishment and comprehensive characterization of a novel dark-reared zebrafish model for myopia studies

**DOI:** 10.1101/2023.11.19.567749

**Authors:** Jiaheng Xie, Patrick T. Goodbourn, Bang V. Bui, Patricia R. Jusuf

**Author notes:** These authors are co-corresponding senior authors (Bang V. Bui, Patricia R. Jusuf).

## Abstract

**Purpose:** Myopia is predicted to impact approximately 5 billion people by 2050, necessitating mechanistic understanding of its development. Myopia results from dysregulated genetic mechanisms of emmetropization, caused by over-exposure to aberrant visual environments; however, these genetic mechanisms remain unclear. Recent human genome-wide association studies have identified a range of novel myopia-risk genes. To facilitate large-scale *in vivo* mechanistic examination of gene-environment interactions, this study aims to establish a myopia model platform that allows efficient environmental and genetic manipulations.

**Methods:** We established an environmental zebrafish myopia model by dark-rearing. Ocular biometrics including relative ocular refraction were quantified using optical coherence tomography images. Spatial vision was assessed using optomotor response (OMR). Retinal function was analyzed via electroretinography (ERG). Myopia-associated molecular contents or distributions were examined using RT-qPCR or immunohistochemistry.

**Results:** Our model produces robust phenotypic changes, showing myopia after 2 weeks of dark-rearing, which were recoverable within 2 weeks after returning animals to normal lighting. 2-week dark-reared zebrafish have reduced spatial-frequency tuning function. ERG showed reduced photoreceptor and bipolar cell function (*a*- and *b*-waves) after only 2 days of dark-rearing, which worsened after 2 weeks of dark-rearing. We also found dark-rearing-induced changes to expression of myopia-risk genes, including *egr1, vegfaa, vegfab, rbp3, gjd2a* and *gjd2b*, inner retinal distribution of EFEMP1, TIMP2 and MMP2, as well as transiently reduced PSD95 density in the inner plexiform layer.

**Conclusions:** Coupled with the gene editing tools available for zebrafish, our environmental myopia model provides an excellent platform for large-scale investigation of gene-environment interactions in myopia development.

## Introduction

Myopia (short-sightedness) is the most common visual disorder in the world. In 2016, myopia was predicted to impact around half of the world population by 2050.^1^ However, gradual shifts away from outdoor activity and increased screen time, including those caused by the response to COVID, may require an upward revision of prevalence projections.^2^ Therefore, the need to understand the mechanisms underlying myopia development and to develop new intervention strategies is more urgent than ever.

Generally, myopia occurs when the eye grows excessively, and thus its optical components (e.g., lens and cornea, etc.) focus the visual input in front of, rather than on the retinal photoreceptor layer, resulting in blurred distance vision. This reflects a failure of emmetropization to match eye size with its optic power during development. This breakdown is precipitated by over-exposure to aberrant visual environments, which alters molecular signaling cascades for normal ocular growth. Strong evidence indicates that the retina contributes to the local regulation of ocular growth without the need for involvement from the brain.^3–5^ However, underlying molecular signaling for myopia development are poorly understood.

Over the past decades, significant effort has resulted in successful identification of a number of important myopia-related genes. Indeed, human genome-wide association studies (GWAS)^6^ and RNA sequencing^7, 8^ continue to grow the number of novel myopia-risk genes. Which of these genes, and how they might mediate aberrant visual environment induced eye growth remains largely unclear. Thus, a vertebrate model that allows efficient genetic manipulation along with reliable and robust induction of environmental myopia is needed for high-throughput, large-scale analysis of gene function *in vivo*. Animals like chicks, guinea pigs, tree shrews, mice, non-human primates and even tilapia have been successfully used in myopia studies, with environmental myopia induction achieved through wearing of diffusors or negative lenses.^9, 10^ However, with the exception of mice, adequately mature genetic editing tools are not available. This has hindered in-depth investigations of gene-environment interactions in myopia development.

The zebrafish (*Danio rerio*) has become a popular laboratory animal model for studies of development, disease modelling, drug screening and neuronal activity, to name a few. Zebrafish have conserved genetics (possessing >80% of disease-associated human orthologues),^11^ and share ocular similarities with humans.^12^ Their high fecundity, small size and ease of maintenance make zebrafish an economic high-throughput model. Importantly, their externally growing embryos are accessible for highly efficient genetic manipulation with a range of well-established gene editing approaches.^13^ Zebrafish have not been widely used for myopia studies. Due to their small size, aquatic living environment and lifelong growth, it is not practical to use common approaches, such as diffusor or negative lens, for environmental myopia induction. To our knowledge, only one environmental zebrafish myopia model has been previously reported; the authors reared zebrafish from 5 days post-fertilization (dpf) under a constant dark environment and reported myopia development from 4 weeks post-fertilization (wpf).^14^ However, no further studies have used this model, possibly because of the risk of reduced survival likely from changes in feeding behavior from an very early age.^15^

Here, we established a stable, novel zebrafish myopia model with an optimized dark-rearing protocol. Our comprehensive characterization demonstrates aberrant ocular growth, impaired spatial vision and altered retinal function consistent with myopia. These structural and functional alterations were associated with changes to molecules previously reported in myopia. Combined with the genetic advantages of zebrafish, this novel environmental myopia model has the potential to be a platform for high-throughput in-depth *in vivo* investigation of gene-environment interactions for myopia development.

## Methods

### Animal husbandry

Zebrafish (*Danio rerio*; wild-type AB strains) were maintained and bred in the Fish Facility at the University of Melbourne according to local animal guidelines. Embryos and larvae (prior to sex determination) were grown in Petri dishes in an incubator at 28.5°C up to 5 days post-fertilization (dpf), then introduced to tanks and raised in flow-through systems at 28°C up to 4 weeks post-fertilization (wpf) for experiments.

For dark-rearing, zebrafish tanks were wrapped using black cloth tape (Model number 66623336603; Bear brand, Saint-Gobain Abrasives, Somerton, VIC, Australia) to block light. Zebrafish were induced to blackout tanks at 4 wpf and reared in darkness for 2 days, 2 or 4 weeks. Zebrafish health and survival were monitored daily using dim red light (LED; 17.4 cd.m^-2^, λ_max_ 600 nm).

For the recovery experiment, zebrafish were dark-reared for 2 weeks, which was followed by 0 (baseline of recovery), 1 and 2 weeks under a normal light/dark cycle.

All procedures were performed according to the provisions of the Australian National Health and Medical Research Council code of practice for the care and use of animals following approval from the Faculty of Science Animal Ethics Committee at the University of Melbourne (Project No. 10399) and adhered to the ARVO Statement for the Use of Animals in Ophthalmic and Vision Research.

### Optical coherence tomography (OCT)

OCT was performed immediately after zebrafish were humanely killed using 1000 ppm AQUI-S (#106036, Primo Aquaculture, Narangba, QLD, Australia) in E3 medium. Zebrafish were positioned on a square (2 × 2 cm) of paper towel and transferred onto a moistened polyvinyl alcohol (PVA) sponge fitted snugly in a 35-mm Petri dish. All OCT images were recorded using a spectral domain OCT/OCTA system (Spectralis OCT2; Heidelberg Engineering, Heidelberg, Germany) with the aid of a Digital High Mag lens (78D, Volk Optical Inc., Mentor, Ohio, USA).

The Petri dish was then affixed onto a platform, allowing the zebrafish eye to be aligned with the lens. Surface tension between the moistened sponge/paper towel and the fish kept it in place. Images were acquired in the dorsal-ventral direction with a volume scan pattern of 15° × 5° (2.7 × 0.9 mm). Each volume consisted of 128 B-scans (5 repeats) each consisting of 512 A-scans. When imaging, the OCT was positioned such that the iris was horizontal and a bright reflection can be seen at the apex of the cornea. This ensured that the image captured was aligned with the central axis of the eye. Both eyes were imaged for each zebrafish.

Only for the 2-week timepoint of the recovery experiments, zebrafish eyes were imaged using a Bioptigen Envisu R2200 spectral domain OCT system with an 18-mm telecentric lens (Bioptigen Inc., Durham, NC), due to temporary unavailability of the Spectralis OCT system. A volume scan of 2 × 2 mm was acquired with a depth of 1.7 mm (1000 A-scans per B-scans, 100 horizontal B-scans evenly spaced in the vertical dimension).

Image quantification was performed using a custom MATLAB script. As illustrated in Fig. 1A, axial length is measured as the distance from the corneal apex to the retinal pigment epithelium (RPE). Lens radius is half of the distance from the anterior to posterior lens surface. The retinal radius is the distance from the center of the lens to the RPE, which can be calculated as the difference between axial length and lens radius. Equatorial diameter is the farthest distance between the dorsal and ventral sides of the sclera. With these parameters, relative ocular refraction is given by the ratio of retinal radius to lens radius, which is also called *Matthiessen’s Ratio*, commonly used for quantifying ocular refraction in aquatic species.^14, 16, 17^ To examine relative ocular refraction in the equatorial dimension, we also calculated the ratio of equatorial diameter to lens radius. The ratio of axial length to equatorial diameter provides an index of overall eye shape. Corneal curvature was quantified using the *Kappa* plugin in ImageJ (National Institutes of Health, Bethesda, MD, USA).^18, 19^

**Figure 1.**
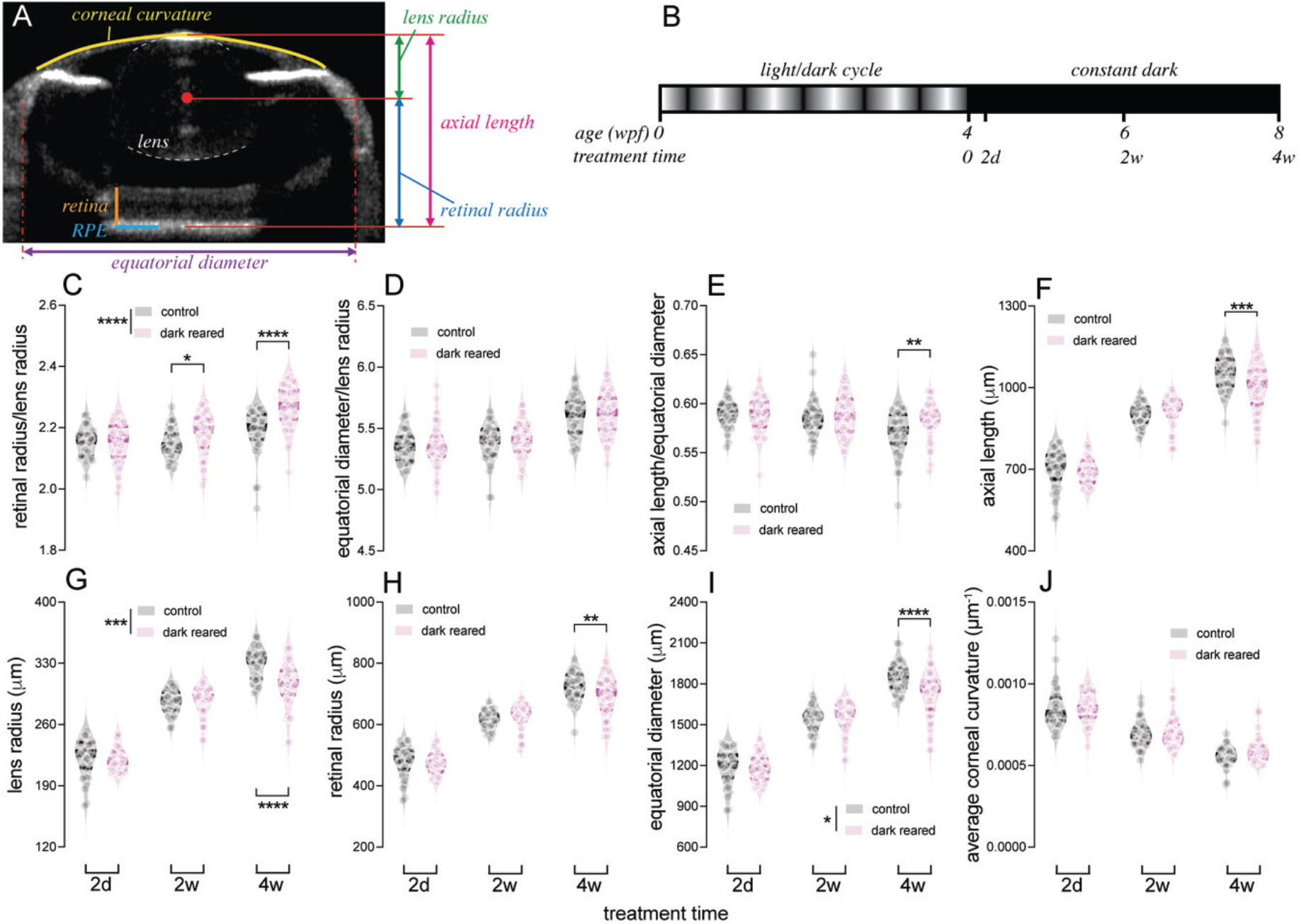
Comparison of ocular biometric parameters between control and dark-reared zebrafish. (**A**) Using optical coherence tomography (OCT) images of zebrafish eyes, ocular parameters including axial length, lens radius, retinal radius, equatorial diameter and corneal curvature were quantified. Lens (white dashed lines), retina (orange solid line) and retinal pigmented epithelium (RPE; light blue solid line) are highlighted in the representative image. (**B**) For dark-rearing, zebrafish were reared under a normal light/dark cycle until 4 weeks post-fertilization (wpf), followed by 2 days (2d), 2 weeks (2w) or 4 weeks (4w) of exposure to a constant dark environment. Controls were age-matched fish reared under normal lighting. (**C**–**J**) To analyze the impact of dark rearing, ratios of (**C**) retinal radius to lens radius, (**D**) equatorial diameter to lens radius, and (**E**) axial length to equatorial diameter, as well as (**F**) axial length, (**G**) lens radius, (**H**) retinal radius, (**I**) equatorial diameter and (**J**) averaged corneal curvature were compared between groups for each dark-rearing duration. In violin plots, dots show data for each individual retina. Thick bars indicate the median, and thin lines indicate interquartile ranges. *N* = 36, 40 and 40 fish per group for 2d, 2w and 4w of dark rearing, respectively. *Two*-way ANOVA was performed. \**P* < *0.05;* \*\**P* < *0.01;* \*\*\**P* < *0.001;* \*\*\*\**P* < *0.0001.* Asterisks next to the vertical bars indicate the group effect, those above or below the horizontal bracket indicate Fisher’s LSD *post-hoc* comparisons.

### Quantification of zebrafish weight

Normal or dark-reared zebrafish were humanely killed using 1000 ppm AQUI-S and weighed using an analytical balance (GR-200, Hurst Scientifc, Perth, Western Australia, Australia).

### Optomotor response (OMR)

#### Apparatus and procedure

The OMR apparatus was adapted from that previously described.^20–22^ A Power Mac G5 computer (Apple Computer, Inc., Cupertino, CA, USA) ran MATLAB 2016b (MathWorks, Natick, MA, USA) with Psychtoolbox extensions.^23^ Stimuli were processed on an ATI Radeon HD 5770 graphics card, with the output sent to a BITS++ video processor (Cambridge Research Systems, Rochester, UK) for increased contrast resolution for stimuli displayed on a M992 flat-screen cathode ray tube (CRT) monitor (Dell Computer Corporation, Round Rock, TX, USA) with its screen facing upwards. During experiments, a zebrafish was placed in custom annulus swimming chamber (inner and outer walls were 46.5 and 69 mm from the center, respectively) with a transparent base, positioned 37.5 mm above the screen. A C922 Pro Stream webcam (Logitech Company, Lausanne, Switzerland; 1080p at 30 Hz), placed 260 mm above the swimming chamber base, controlled using MATLAB recorded videos of each trial for *post-hoc* analysis of zebrafish angular movement.

During a trial, a test stimulus was displayed on the CRT screen below the swimming chamber, rotating at 0.5 rad/s for 30s. Test stimuli were windmill sinusoidal gratings with central spatial frequencies of 0.0086, 0.0171, 0.0343, 0.0685, 0.137 and 0.274 cycle per degree (c/°). A blank grey screen was presented between test stimuli. Test stimuli were generated using the green and blue channels of the CRT monitor with constant red luminance across the screen. A long pass filter (cut-off wavelength 600 nm) was a fixed to the camera to filter out blue and green light, allowing only red light to pass through, thus eliminating the moving stimulus and allowing better visualization of the fish or analysis.

OMR was measured in age-matched controls and 2-week dark-reared zebrafish, after acclimating fish to the swimming chamber for 5 minutes. All experiments were conducted between 9:00 AM and 7:00 PM. After experiments, fish were humanely killed using 1000 ppm AQUIS (#106036, Primo Aquaculture, Narangba QLD, Australia).

#### Video and statistical analysis

The position of zebrafish was tracked using a published Python package *Stytra*.^24^ Using custom MATLAB algorithms, we calculated the overall angular movement of zebrafish in the direction of the rotating grating for each trial as the optomotor index (OMI). Normalized OMIs for spatial-frequency tuning functions were calculated by normalizing OMI data to the averaged OMI of control fish for 0.0343 c/° (i.e., the condition with largest response). Spatial-frequency tuning functions were fit using a log-Gaussian function using a least-squares criterion, to return amplitude (i.e., height of the peak), peak spatial frequency (i.e., spatial frequency at which amplitude peaked), and bandwidth (i.e., standard deviation).

### Electroretinography (ERG)

Scotopic ERGs were measured for age-matched controls and 2 days or 2 weeks dark-reared fish in a lightproof room, as adapted from our published methods.^25^ Fish were dark adapted (>8 hours, overnight) prior to experiments. All procedures were conducted under dim red illumination (17.4 cd·m^−2^; λ_max_ = 600 nm). For recording, a fish was humanely killed by immersing in 0.1% tricaine MS-222 in E3 medium. When gills had stopped moving and fish were unresponsive to a gentle touch, the head was removed and immediately rinsed in E3 medium. The eyes were quickly dissected using fine-tipped tweezers under a dissecting microscope. A small incision was made around the central cornea for each eye to allow a small amount of aqueous humor to flow out. This helps keep the recording electrode to moist, increasing conductivity and minimizing noise. The dissected eyes were moistened with multiple drops of E3 medium using a 3-mm Pasteur pipette. The eyes were then transferred onto a square of paper towel (2 × 2 cm) on a moist PVA sponge platform in a Faraday cage. Both eyes were measured at the same time. Under a microscope, the recoding electrodes (99% silver, chloride-electroplated; 0.3 mm thickness) were positioned to gently touch the central apex, and the reference electrodes were inserted into the sponge platform. After electrode placement, the Ganzfeld bowl was moved to cover the platform, and the eyes were allowed to dark adapt for 3 minutes. ERG responses were recorded with flash stimuli of −2.11, −0.81, 0.72, 1.89, or 2.48 log cd·s·m^−2^. At −2.11 and −0.81 log cd·s·m^−2^, three repeats were measured with an inter-flash interval of 10 seconds. At 0.72 to 2.48 log cd·s·m^−2^, a single response was measured with 60 seconds between flashes. All experiments were performed between 9:00 AM and 7:00 PM at room temperature. Amplitudes of the *a*- and *b*-waves were measured from baseline to the negative *a*-wave trough and from the *a*-wave trough to the *b*-wave peak, respectively. Implicit times of the *a*- and *b*-waves were measured from stimulus onset to the *a*-wave trough and the b-wave peak, respectively.

### Locomotor response

Age-matched control and 2-week dark-reared zebrafish were transferred to a 6-well plate (well diameter 34.8 mm) with one fish per well. Well plates were positioned on a monitor screen facing upwards and fish were allowed to acclimatize for 10 minutes without any stimulus on the screen. A C922 Pro Stream webcam was fixed 297.5 mm above the screen to take videos of zebrafish swimming. During the test, the monitor displayed a white screen (bright luminance 100.8 cd.m^-2^) or a grey screen (dim luminance 50.4 cd.m^-2^), videos were taken for 5 minutes at each luminance level. Videos were processed using a Python package *Stytra*^24^ to return zebrafish positions, allowing total swimming distance and averaged swimming speed to be calculated (MATLAB).

### Gene expression analysis

Zebrafish were humanely killed by 1000 ppm AQUIS (#106036, Primo Aquaculture, Narangba QLD, Australia) and stored in RNALater (AM7020, Thermo Fisher, Waltham, MA, USA) at – 20°C if not immediately used for RNA isolation. For RNA isolation, eyes were dissected from zebrafish and lenses were removed. For each sample, the total RNA of four eyes from two fish was isolated using TRIzol (or TRI-reagent; AM9738, Thermo Fisher, Waltham, MA, US)^27^ and quantified using a Nanodrop 1000 (Thermo Fisher Scientific, Waltham, MA, USA). 1 ng of the total RNA was reverse-transcribed into cDNA using a Tetro cDNA synthesis kit (BIO-65042, Bioline, London, UK). Quantitative PCR (RT-qPCR) assays were performed on a CFX96 real-time PCR system (Bio-Rad, Hercules, CA, USA) using SsoAdvanced Universal SYBR Green Supermix (12 μL final mix per reaction; 1725272, Bio-Rad). In this study, mRNA levels of *egr1*, *efemp1*, *tgfb1a*, *tgfb1b*, *vegfaa*, *vegfab*, *igf1*, *fgf2*, *wnt1b*, *rbp3*, *gjd2a* and *gjd2b* were quantified. Primers were designed using Primer Premier 5.0^28^ as listed in Table 1. A housekeeping gene *elongation factor 1-alpha* (*ef1a*) served as an internal control.^26^ For each gene, there were 4 samples per group and 3 replicates per sample. Calculation of expression levels was performed using the 2^−ΔΔCT^ method^29^ and the transcript levels were normalized to the *ef1a* transcript levels in control eyes at each time point.

**Table 1.**
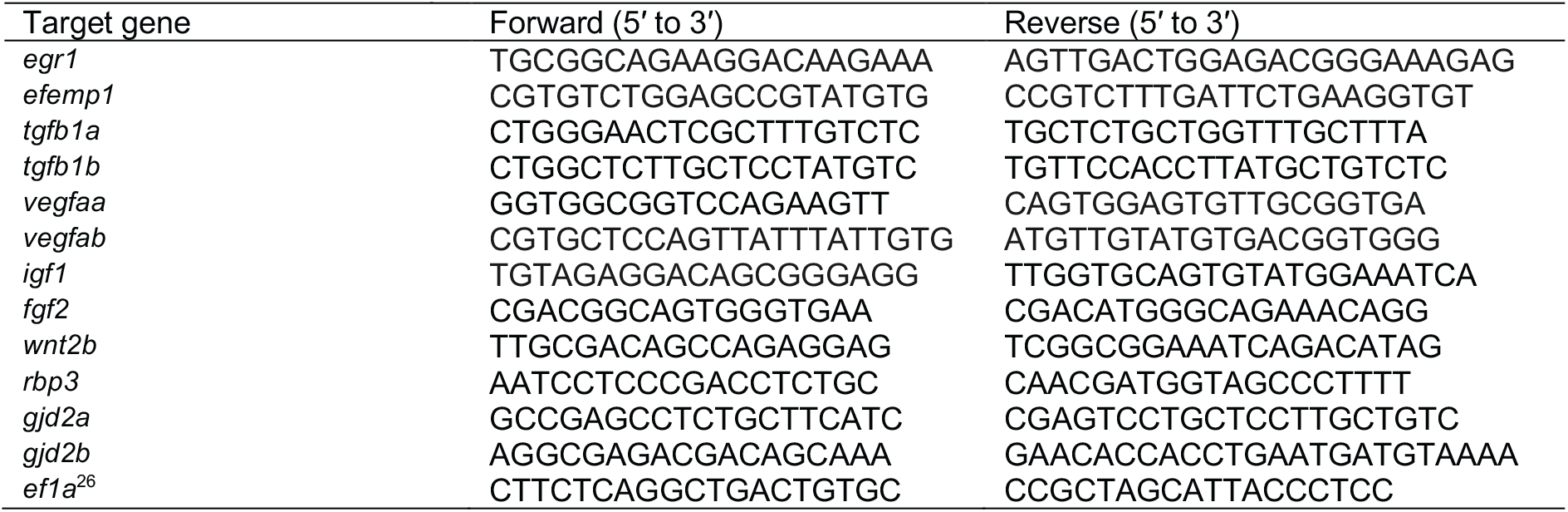
Primers used for RT-qPCR.

### Histology

#### Immunohistochemistry

Dissected zebrafish eyes were fixed in 4% paraformaldehyde (PFA) in PBS for 48 hours at 4°C. They were cryoprotected in 30% sucrose in PBS, embedded in Tissue-Tek OCT compound, and cryosectioned (20 μm; CM 1860 Cryostat; Leica, Wetzlar, Germany). Antibody staining was carried out at room temperature using standard protocols. Slides were blocked in 5% fetal bovine serum (FBS) for 30 minutes and incubated overnight in rabbit anti-EFEMP1 (ARP41450_P050, 1:600; Sapphire Bioscience, NSW, Australia), sheep anti-TIMP2 (1:1000; kind gift from the Itoh Lab in University of Oxford), rabbit anti-MMP2 (AS-55111, 1:250; AnaSpec, Fremont, CA, USA), mouse anti-TH (1:1000, MAB318; Merck, Rahway, NJ, United States) or rabbit anti-PSD-95 (ab18258, 1:100; Abcam, Cambridge, UK) primary antibodies diluted in FBS. Slides were subsequently incubated for 2 hours in secondary antibodies (all 1:500; Thermo Fisher Scientific, Waltham, MA, USA) diluted in 5% FBS. The secondary antibodies used were goat anti-rabbit Alexa Fluor 488 (A11008), chicken anti-rabbit Alexa Fluor 647 (A21443), donkey anti-sheep Alexa Fluor 488 (A11015), goat anti-mouse Alexa Fluor 488 (A11001). Nuclei were counterstained with 4′, 6-diamidino-2-phenylindole (DAPI; D9542-10MG, 1:10000; Sigma-Aldrich) in PBS for 20 minutes, and sections were mounted in Mowiol (81381-250G; Sigma-Aldrich). Stained sections were imaged at a distance of <100 µm from the center of the optic nerve, using a Nikon A1R confocal microscope (Nikon, Minato City, Tokyo, Japan) with a 40× air objective lens (NA 0.95). For all confocal imaging, the deconvolution function of the NIS elements software (AR4.6, Nikon, Minato City, Tokyo, Japan) was applied to minimize background noise.

#### Image analysis

To analyze the relative distribution of EFEMP1, TIMP2 and MMP2, images were pre-processed in FIJI.^19^ In each image, an inner retinal region was selected by drawing a line that followed the contour of the inner plexiform layer (IPL) with a line thickness of 1000 pixel, which includes the inner nuclear layer (INL; including the amacrine cell layer, ACL), IPL and ganglion cell layer (GCL). This line allowed the retinal cross section to be straightened using the ‘*Straighten*’ function.

Straightened images were rotated, positioning the INL on the left and GCL on the right. The widths of the straightened images were resized using Adobe Photoshop to normalize IPL thickness to the averaged IPL thickness of the control group relevant to that time point. Regions of interest (ROI) measuring 500-pixel width and heights of 415 (EFEMP1-stained 2-day-treated retinae), 440 (EFEMP1-stained 2-week-treated retinae) or 450 pixels (all TIMP2- and MMP2-stained images) were used to return expression levels using the ‘*Plot Profile*’ function in FIJI. Brightness profiles across the ACL, IPL and GCL were normalized to the highest value. To examine relative change in the three layers, normalized intensity was summed throughout the ACL (pixel 1–90), IPL (2-day: pixel 91–273 for EFEMP1, pixel 91– 310 for TIMP2 and MMP2; 2-week: pixel 91–310 for EFEMP1, pixel 91–350 data points for TIMP2 and MMP2) and GCL (2-day: pixel 274–415 for EFEMP1, pixel 311–450 for TIMP2 and MMP2; 2-week: pixel 311–440 for EFEMP1, pixel 351–450 data points for TIMP2 and MMP2) for each image. Please note that at each time point, the range for EFEMP1 was different from that of TIMP2 or MMP2. This is likely because fish used for EFEMP1 histological analysis were from a different batch from those used for TIMP2 and MMP2 analysis and thus the thickness of the IPL was different between batches.

For analysis of dopaminergic cell number, TH positive cells were manually counted for each image in a masked fashion. For quantification of PSD95 puncta in the IPL, a region of interest (ROIs; 50 × 20 μm) were randomly selected from the IPL approximately in the central retinal region using FIJI. For each ROI, thresholding using the ‘*Mean*’ method was performed^30^ with the ‘*Invert*’ lookup table (LUT) function to select areas containing no puncta to create a background mask. All potential puncta in the ROI were identified as dots using the ‘*Find Maxima*’ function (prominence = 3) to output a ‘*single points*’ image. Dots identified in the background area (i.e., noise) were masked using the ‘AND’ algorithm of the ‘*Image Calculator*’ function with the ‘*single points*’ image and the background mask. The remaining dots in the ROI were considered true puncta and quantified for analysis. For all histological analyses, one section per retina was used and for each fish both eyes were used.

### Statistical analysis

Except for OMR, all statistical analyses were performed using Prism 9 (GraphPad, San Diego, CA, USA). Data were plotted as the mean ± SEM or median with interquartile ranges as indicated in figure legends. Data were compared either using a two-way ANOVA with Fisher’s LSD *post-hoc* multiple comparisons or two-tailed unpair *t-*test. Statistical comparisons of OMR were performed using MATLAB. Data were plotted as the mean ± SEM or mean with 95% confidence intervals as indicated in figure legends. To test whether spatial-frequency tuning functions differed between groups, an omnibus *F*-test was used to compare the goodness of fit (*r*^2^) of a full model, in which parameter estimates of each group could vary independently, with that of a restricted model, in which parameters were constrained to be the same across groups. To determine whether specific parameter estimates differed between groups, a nested *F*-test was used to compare a full model with a restricted model in which one parameter was constrained to be the same across groups.^31^ For all analyses, a criterion of α = 0.05 was used to determine significance. Precise sample sizes are indicated in the figure legends.

### Data availability

Data for the present study are available on the Open Science Framework (https://osf.io/kzq34/).

## Results

### Dark-rearing induces axial myopia in zebrafish

In this study, we used a novel dark-rearing protocol to induce myopia in zebrafish and measured ocular biometrics using OCT B-scan images (Fig. 1A). Fish were first reared under normal light/dark cycles until 4 weeks of age, followed by 2 days, 2 or 4 weeks of dark-rearing (Fig. 1B). Control groups were age-matched fish reared under normal lighting. Using the retinal radius to lens radius ratio, we found that dark-rearing resulted in axial myopia in zebrafish eyes. The difference between groups was significant from 2 weeks post-treatment (*P* = *0.0302*) and increased with treatment duration (*P* < *0.0001* for 4-week treated fish; treatment × time: *P* = *0.0007*, 2-way ANOVA; Fig.1C). We did not find a myopic shift in the equatorial dimension in this experiment (Fig. 1D). The ratio of axial length to equatorial diameter for 4-week dark-reared fish was significantly higher than that of the control group (*P* = *0.0035*), suggesting that axial elongation resulted in an eye shape change (Fig. 1E). We observed that although increasing dark-rearing duration increased the magnitude of myopia induction, it may also be associated maldevelopment; 4-week dark-reared zebrafish showed reduced axial length (*P* = *0.0004*; Fig. 1F), lens radius (*P* < *0.0001*; Fig. 1G), retinal radius (*P* = *0.0058*; Fig. 1H) and equatorial diameter (*P* < *0.0001*; Fig. 1I), compared to control fish.

In addition, we examined the effect of dark-rearing on general development for zebrafish by quantifying body weight. Our results suggested that dark-rearing had an effect on general development (*P* = *0.0004*, 2-way ANOVA), but between-group differences were only significant after 4 weeks of dark-rearing (*P* = *0.0005*; Fig. 2). The average corneal curvature in zebrafish was not affected by dark-rearing (Fig. 1J). It is important to mention that we did not encounter any survival issues in this dark-reared model, which had over 90% survival rate in all our experiments (Table 2). Taken together, our novel dark-reared zebrafish model showed axial myopia after 2 weeks of dark-rearing, but with longer exposure to constant dark, maldevelopment can occur. Therefore, in the subsequent experiments, we focused on the 2-week dark-reared model.

**Figure 2.**
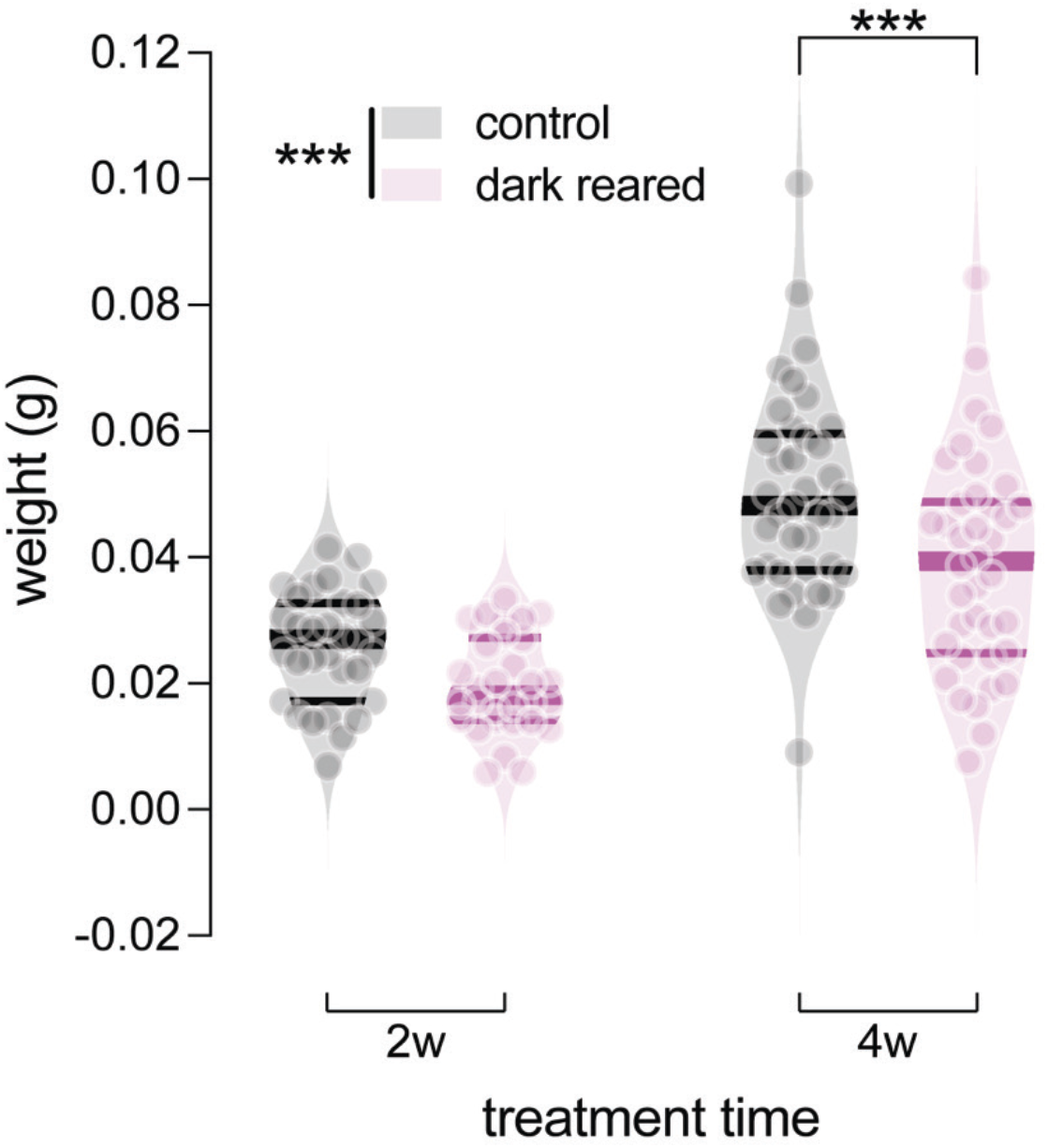
Comparison of body weight between control and dark-reared zebrafish at 2 weeks (2w) and 4 weeks (4w) post-treatment. For 2-week treated groups, there were 31 control and 30 dark-reared fish. For 4-week treated groups, there were 37 control and 38 dark-reared fish. Violin plots show data for individual retinae. Thick bars indicate medians, and thin lines indicate interquartile ranges. *Two*-way ANOVA was performed. \*\*\**P* < *0.001.* Asterisks next to the vertical bars indicate the group effect, those above the horizontal bracket indicate Fisher’s LSD *post-hoc* comparisons.

**Table 2.** Survival of zebrafish after 4 weeks of dark-rearing experiments.

| experiments | 1 | 2 | 3 | 4 | 5 | 6 | average |
| --- | --- | --- | --- | --- | --- | --- | --- |
| survival | 93.18%<br>(41/44) | 94.44%<br>(34/36) | 97.73%<br>(43/44) | 100%<br>(50/50) | 100%<br>(36/36) | 90.72%<br>(88/97) | 96.01% |

### Dark-reared zebrafish recover from myopia after returning to normal lighting

Animal myopia models, such as chicks, monkeys and tree shrews,^32–34^ can recovery from form-deprivation or lens-induced myopia, providing the potential to investigate mechanisms that can reverse myopia development. Here, we tested whether our zebrafish myopia model was able to recover from dark-rearing-induced myopia. For this, zebrafish were reared under normal light-dark cycles until 4 weeks post-fertilization (wpf), followed by 2 weeks of dark-rearing before being returned to normal lighting to recovery (Fig. 3A). The control groups were age-matched fish reared under a normal light/dark cycle for the same duration. Our results showed that the retinal radius to lens radius ratio was significantly higher in 2-week dark-reared fish, compared to control fish (*P* = *0.0033*), indicating axial myopia in dark-reared fish. This phenotype disappeared after 1 week of recovery (Fig. 3B). Interestingly, in this experiment we also found evidence of equatorial myopia in dark-reared fish (effect of dark-rearing: *P* = *0.0017*, 2-way ANOVA) at baseline (*P* = *0.0177*) and 1 week of recovery (*P* = *0.0251*), an effect was normalized after 2 weeks of recovery (Fig. 3C). It is therefore not surprising that overall there was a flatter eye shape (axial length to equatorial diameter ratio) in dark-reared than in control fish during recovery (*P* = *0.0063*, 2-way ANOVA); there was no different between groups at baseline, but the dark-reared group showed lower ratio at 1 week of recovery (*P* = *0.0081*), likely due to the axial recovery combined with the unchanged equatorial myopia. The significant difference in eye shape disappeared following 2 weeks of recovery (Fig. 3D).

**Figure 3.**
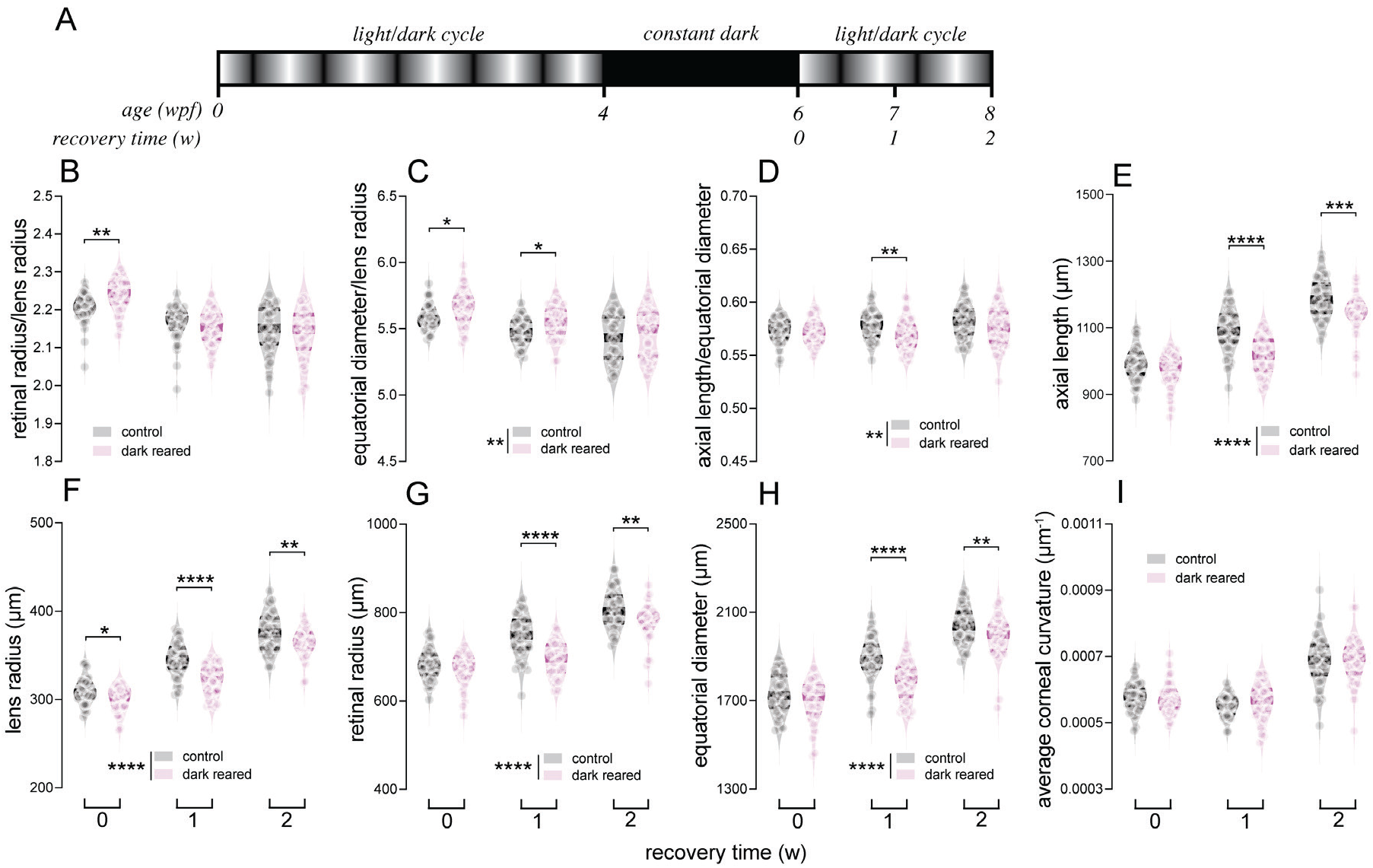
Ocular biometric parameters of control and 2-week dark-reared fish at 0, 1 and 2 weeks (w) post-recovery. (**A**) For rearing treatments, wild-type fish were reared under a constant dark environment for 2 weeks from 4 weeks post-fertilization (wpf), followed by recovery in normal light/dark rearing. (**B**–**I**) Parameters analyzed were (**B**) retinal radius to lens radius ratio, (**C**) equatorial diameter to lens radius ratio and (**D**) axial length to equatorial diameter ratio, as well as (**E**) axial length, (**F**) lens radius, (**G**) retinal radius, (**H**) equatorial diameter and (**I**) averaged corneal curvature, were compared between control and treated fish at 0 (baseline), 1 and 2 weeks of recovery. Violin plots show data for individual retinae with thick bars indicating medians, and thin lines indicating interquartile ranges. *N* = 40 control fish for each timepoint. *N* = 38, 40 and 37 dark-reared fish for 0, 1 and 2 w of recovery, respectively. *Two*-way ANOVA was performed. \**P* < *0.05;* \*\**P* < *0.01;* \*\*\**P* < *0.001;* \*\*\*\**P* < *0.0001.* Asterisks next to the vertical bars indicate the group effect, those above the horizontal bracket indicate Fisher’s LSD *post-hoc* comparisons.

Other parameters including the axial length (Fig. 3E), lens radius (Fig. 3F), retinal radius (Fig. 3G) and equatorial diameter (Fig; 3H), were lower in dark-reared fish compared with control fish during the 2-week period of recovery (effect of treatment: *P* < *0.0001* for all, 2-way ANOVA). There were no differences, except for lens radius (*P* = *0.0150*), between the groups at the baseline measurement. Differences in these parameters between groups became highly significant after 1-week recovery (*P* < *0.0001* for all), followed by a drop of the significance (*P* = *0.0006*, *0.0015*, *0.0014* and *0.0074*, respectively). Therefore, although the effect of dark-rearing on ocular growth might remain for a short period of time (i.e. 1 week), our data here suggested that fish dark-reared for 2-weeks should fully recover after 2 or more weeks after returning to a normal light environment. Again, the average corneal curvature was not different between groups after dark-rearing and during recovery from myopia (Fig. 3I).

### Altered visual behavior in dark-reared myopic zebrafish

Next we considered whether axial myopia in 2-week dark-reared zebrafish manifest changes in visual behavior using optomotor responses (OMR). Using an omnibus *F*-test, we showed that spatial-frequency tuning functions significantly differed between control and 2-week dark-reared zebrafish (*P* = *0.0145*; Fig. 4A). Comparing parameters, dark-reared fish showed a lower function amplitude than control fish (*P* = *0.0155*; Fig. 4B), but there was no difference in function peak frequency (Fig. 4C) or bandwidth (Fig. 4D). These results indicated impaired spatial vision at the behavioral level in dark-reared zebrafish.

**Figure 4.**
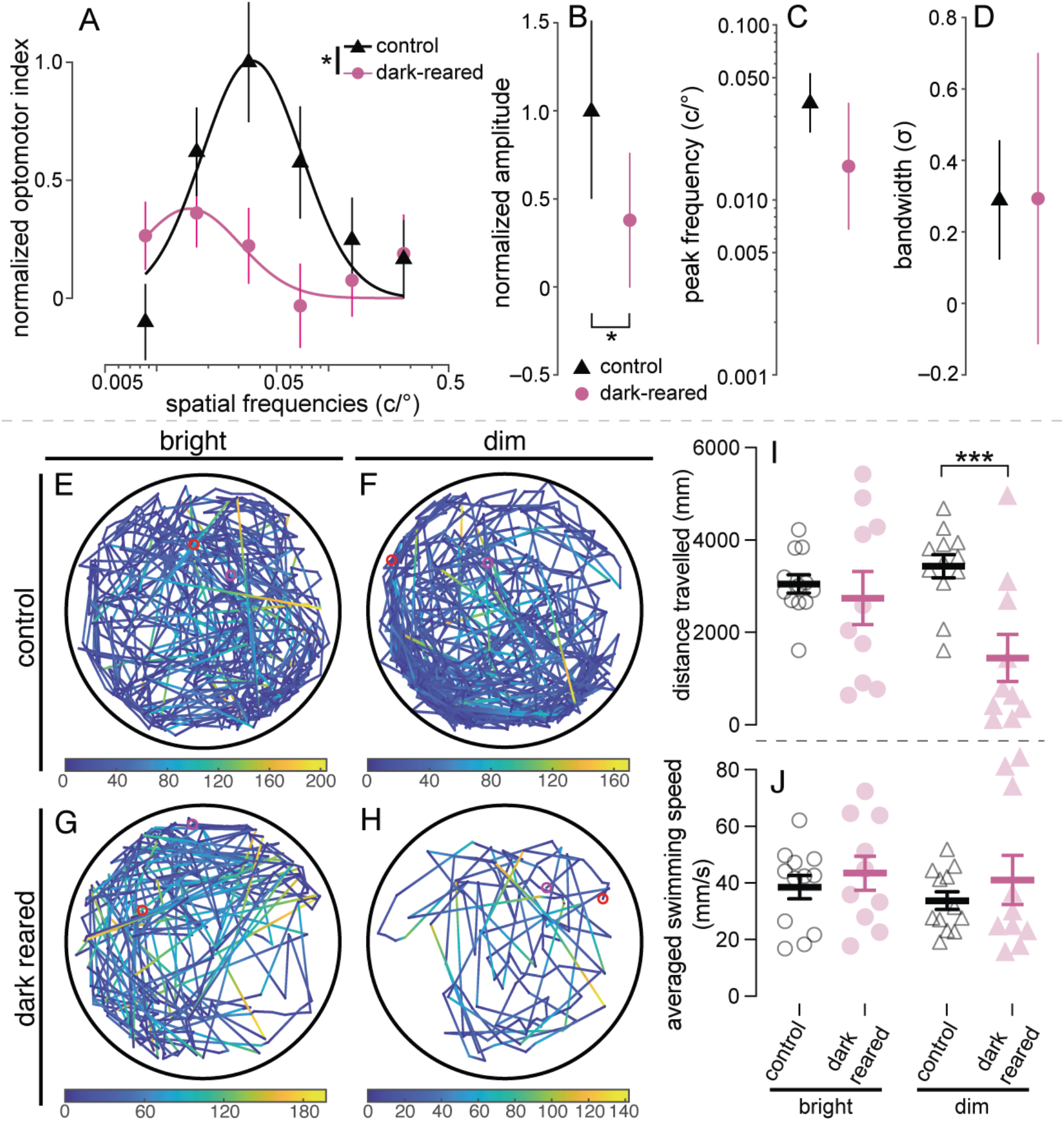
Assessment of visual behavior for control and 2-week dark-reared fish. (**A**–**D**) Spatial-frequency tuning functions were measured using optomotor responses (OMR) in control (*N* = 31) and 2-week dark-reared zebrafish (*N* = 30). (**A**) Spatial-frequency tuning functions are three-parameter log-Gaussian functions fit to the data by minimizing the least-square error. Error bars show ± SEM. The fitted parameters, including (**B**) normalized amplitude, (**C**) peak frequency and (**D**) bandwidth, were compared between groups. Error bars indicate 95% confidence intervals. Nested *F*-test was performed. (**E**–**J**) Locomotor responses of control and 2-week dark-reared zebrafish. (**E**–**H**) Representative 5-minute swimming traces for control fish (*N* = 12) under bright and dim luminance (**E** and **F**, respectively), as well as dark-reared fish (*N* = 10) under bright and dim luminance (**G** and **H**, respectively). In each trace, the red and magenta circles indicate start and end positions, respectively. Swimming speeds during recording are color coded onto swimming traces as per the colormaps below each trace. Black circles around traces indicate the wells that contained the fish. Total distance travelled (**I**) and averaged swimming speeds (**J**) were quantified. *Two*-way ANOVA with Fisher’s LSD was performed. For all statistics, \**P* < *0.05*; \*\*\**P* < *0.001*. Asterisks next to the vertical bars indicate the group effect, those above or below the horizontal bracket indicate *post-hoc* comparisons.

Although our quantification of the zebrafish weight at 2 weeks post-treatment did not show significant difference between groups (Fig. 2), one may still be concerned that dark-rearing can lead to systemic effects, and therefore reduced optomotor responses in dark-reared fish may represent deficits of overall development rather than vision. For this, we performed locomotor tests to compare spontaneous swimming between control and 2-week dark-reared zebrafish. We found that there was no difference between groups in total swimming distance under bright luminance (Fig. 4E–F and 4I), while dark-reared zebrafish swum significantly less than age-matched controls under dim luminance (*P* = *0.0009*; Fig. 4G–H). There was no difference in the average swimming speed between groups under both lighting conditions (Fig. 4J). These results suggested that dark-reared zebrafish were able to swim as well as controls if they were stimulated with visual stimuli. The reduction in spontaneous swimming in dark-reared fish placed in dim environments is consistent with impaired visual function.

### Altered retinal function in dark-reared myopic zebrafish

As local retinal mechanisms are thought to mediate myopia development arising from aberrant visual input,^3–5^ we sought to quantify changes in retinal function using electroretinography (ERG). After 2 days (Fig. 5A) and 2 weeks (Fig. 5F) of dark-rearing, ERG responses in treated fish were significantly smaller than age-matched controls. At 2 days post-treatment, dark-reared fish had lower ERG photoreceptoral *a*- (effect of treatment: *P* = *0.0391*, 2-way ANOVA; Fig. 5B) and bipolar cell *b*-wave amplitudes (effect of treatment: *P* = *0.0119*, 2-way ANOVA; *P* = *0.0441* at 2.48 log cd.s.m^-2^; Fig. 5D). 2-day dark-rearing also led to a slower *a*-wave implicit time, particularly at the lowest intensity –2.11 log cd.s.m^-2^ tested (*P* = *0.0349*; Fig. 5C). There was no difference between groups for *b*-wave implicit time (Fig. 5E). These differences suggest that retinal signaling is altered, following 2 days of dark-rearing, before the development of myopic ocular structure.

**Figure 5.**
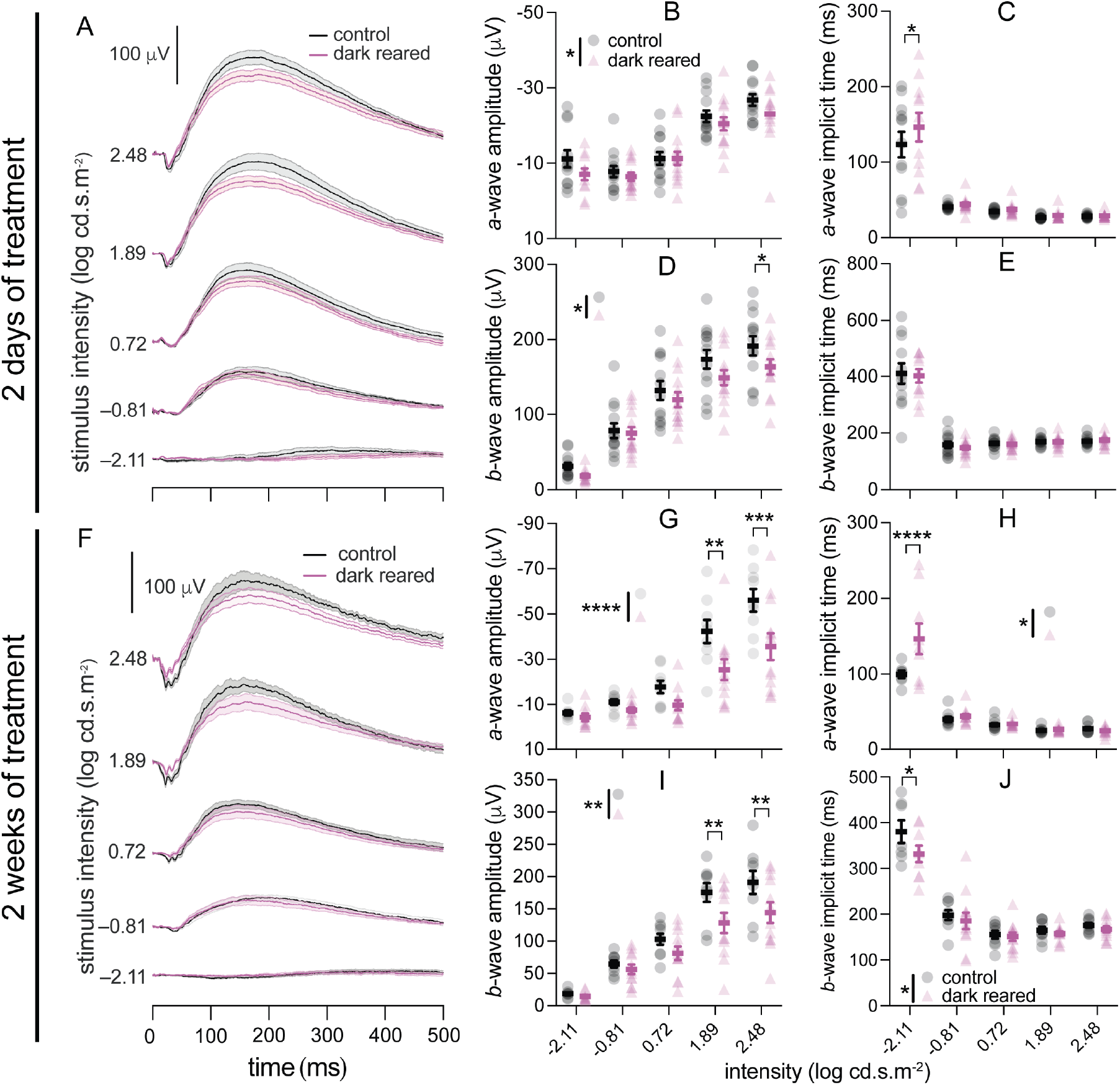
Scotopic electroretinography (ERG) of control and dark-reared fish after (**A**–**E**) 2 days and (**F**–**J**) 2 weeks of treatment. (**A, F**) Averaged ERG traces for control (black) and dark-reared (pink) zebrafish, at −2.11, −0.81, 0.72, 1.89, and 2.48 log cd·s·m^−2^. Scale bar: 100 µV. The light bands around group average traces represent ±1 SEM. The remaining panels show group average (±SEM) *a*-wave amplitude (**B**, **G**), *a*-wave implicit time (**C**, **H**), *b*-wave amplitude (**D**, **I**), and *b*-wave implicit time (**E**, **J**) for control and dark-reared fish. There were 13 control and 14 dark-reared fish for 2 days of treatment, and 9 control and 12 dark-reared fish for 2 weeks of treatment. *Two*-way ANOVA was performed. \**P* < *0.05;* \*\**P* < *0.01;* \*\*\**P* < *0.001;* \*\*\*\**P* < *0.0001.* Asterisks next to the vertical bars indicate the group effect, those above the horizontal bracket indicate Fisher’s LSD *post-hoc* comparisons.

The difference in the ERG were more apparent after 2 weeks of dark-rearing, with a reduction in *a*- (effect of treatment:, *P* < *0.0001*, 2-way ANOVA; *P* = *0.0018* and *P* = *0.0002* for 1.89 and 2.48 log cd.s.m^-2^, respectively; Fig. 5G) and *b*-wave amplitudes (effect of treatment: *P* = *0.0016*, 2-way ANOVA; *P* = *0.0069* and *P* = *0.0072* for 1.89 and 2.48 log cd.s.m^-2^, respectively; Fig. 5I). 2-week dark-reared fish had slower *a*-wave responses compared to control fish (effect of treatment: *P* = *0.0126*, 2-way ANOVA), especially at the lowest tested intensity (*P* < *0.0001* at –2.11 log cd.s.m^-2^; Fig. 5H). Additionally, 2-week dark-reared group also had faster *b*-wave implicit times (effect of treatment: *P* = *0.0362*, 2-way ANOVA) compared with controls, particularly at –2.11 log cd.s.m^-2^ (*P* = *0.0128*; Fig. 5J). A smaller but faster *b*-wave is likely to reflect dark-rearing-induced inner retinal changes, perhaps change to bipolar cells and/or their interactions with amacrine cell-mediated inhibitory pathways.

### Altered expression of myopia-associated genes in dark-reared myopic zebrafish

To investigate potential genetic pathways involved in what could be considered initiating (2-days) and consolidating (2-weeks) phases of myopia development in dark-reared zebrafish, we analyzed expression of myopia-associated genes. *Early growth response 1* (*EGR1*) is a well-known marker for the direction of ocular growth (myopic or hyperopic)^35, 36^ and its absence leads to myopia in mice.^37^ Our results showed that in zebrafish, as expected *egr1* was down-regulated at both 2 days and 2 weeks after dark-rearing (effect of treatment: *P* = *0.0001*, 2-way ANOVA; *P* = *0.0030* and *P* = *0.0017*, respectively, Fisher’s LSD *post-hoc* test; Fig. 6A).

**Figure 6.**
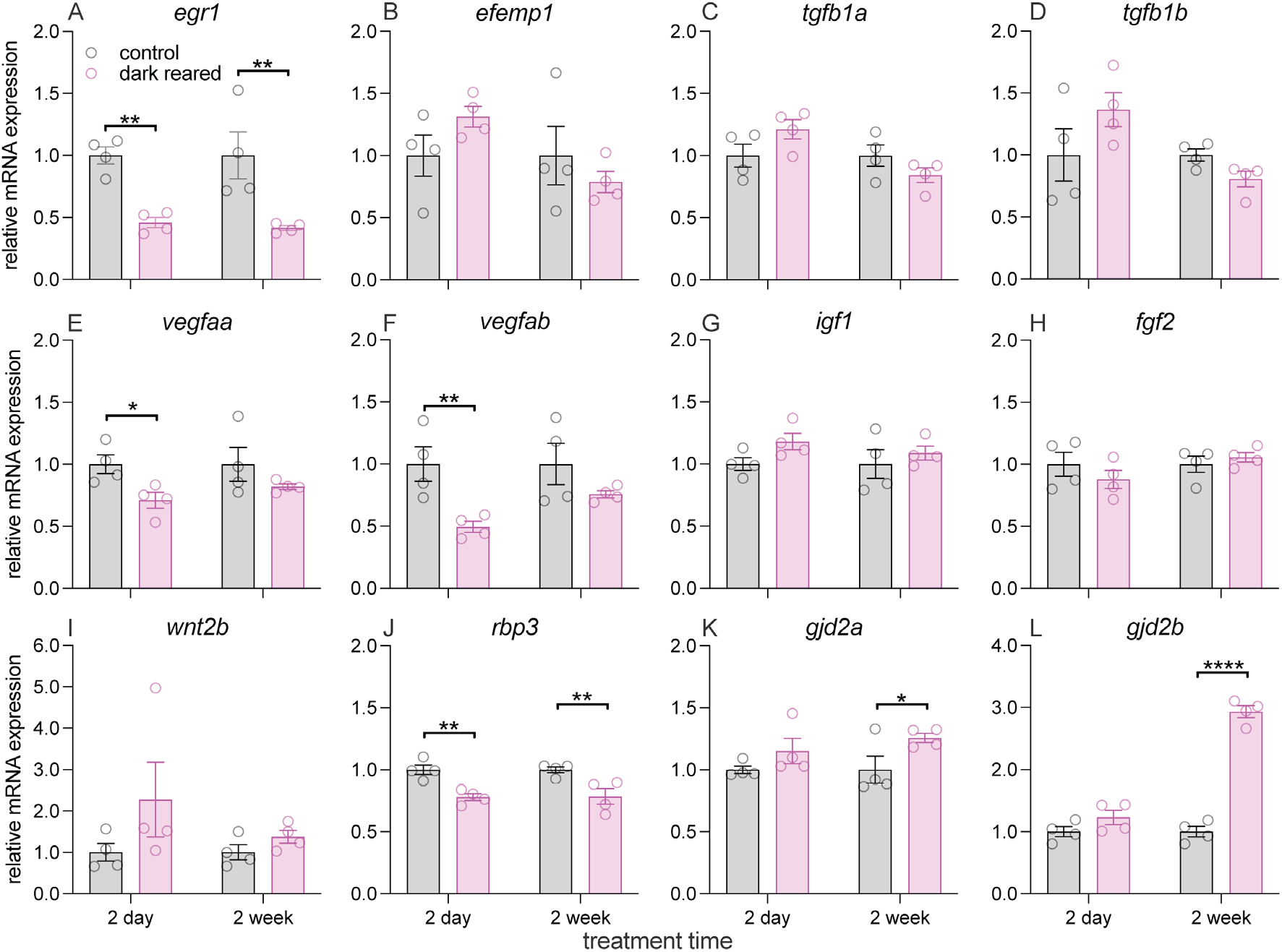
Quantitative analysis of myopia-associated genes in eyes of control and dark-reared zebrafish after 2 days and 2 weeks of treatment. (**A**–**L**) Relative mRNA expression of *egr1* (**A**), *efemp1* (**B**), *tgfb1a* (**C**), *tgfb1b* (**D**), *vegfaa* (**E**), *vegfab* (**F**), *igf1* (**G**), *fgf2* (**H**), *wnt2b* (**I**), *rbp3* (**J**), *gjd2a* (**K**) and *gjd2b* (**L**) were examined using quantitative PCR (RT-qPCR). Error bars show ± SEM. *N* = 4 per group. *Two*-way ANOVA was performed. \**P* < *0.05*; \*\**P* < *0.01*; \*\*\*\**P* < *0.0001*. Asterisks above the horizontal bracket indicate Fisher’s LSD *post-hoc* comparisons.

*EGF containing fibulin extracellular matrix protein 1* (*EFEMP1*) has recently been associated with refractive error in a human genome-wide association study (GWAS)^6^ and mutations of this gene in humans can result in a number of ocular conditions, including Malattia Leventinese, Doyne honeycomb retinal dystrophy,^38^ familial juvenile-onset open-angle glaucoma^39^ and progressive myopia.^40^ However, we did not find a change in *efemp1* expression in dark-reared zebrafish (Fig. 6B).

*Transforming growth factor beta 1* (*TEGFB*) has also been linked to high myopia in a recent meta-analysis of human data.^41^ However, our data showed no transcriptional changes for the two zebrafish analogues of this gene (*tgfb1a* and *tgfb1b*) following dark-rearing (Fig. 6C–D).

Retinal Vascular Endothelial Growth Factor-A (VEGFA) concentration has been negatively correlated with myopia severity in marmosets^42^ and its inhibition enhanced the effect of form-deprivation myopia in guinea pigs.^43^ Here, we showed that both zebrafish analogues of this gene (*vegfaa* and *vegfab*) were, as expected, down-regulated (effect of treatment: *P* = *0.0173*, and *P* = *0.0054*, respectively, 2-way ANOVA), especially at the initiating stage of myopia development (*P* = *0.0330* and *P* = *0.0072*, respectively), but expression was similar between groups after 2 weeks of dark-rearing (Fig. 6E–F).

Single-nucleotide polymorphisms (SNPs) rs2162679 of *insulin-like growth factor 1* (*IGF1*) has been associated with myopia in Chinese children.^44^ On the other hand, with form-deprivation myopia, *fibroblast growth factor 2* (*FGF2*) was found to be down-regulated in the chick sclera^45^ but up-regulated in the guinea pig retina during myopia development.^46^ Injection of a combination of IGF1 and FGF2 caused extreme myopia in chicks,^47^ again implicating these two growth factors in regulation of ocular growth. Our results did not show mRNA level changes for these genes in dark-reared zebrafish (Fig. 6G–H).

Although up-regulation of retinal *wnt2b* has been observed in a murine model of form-deprivation myopia,^48^ our data shows that dark-rearing-induced myopia in zebrafish was not associated with transcriptional changes in *wnt2b* (Fig. 6I).

Knockout of the interphotoreceptor retinoid-binding protein (IRBP; or Rbp3 in zebrafish) in mice leads to excessive enlargement of the eye, indicating that in addition to its role in photopigment recycling, the *irbp* gene is also important for regulating ocular growth.^49^ After dark-rearing, zebrafish eyes showed significantly lower *rbp3* expression (effect of treatment: *P* = *0.0002*, 2-way ANOVA; *P* = *0.0027* and *P* = *0.0032* for 2 days and 2 weeks post-treatment, respectively; Fig. 6J), implicating involvement of an altered visual cycle in myopia development in this model.

Last but not least, the *GJD2* gene (*gjd2a* and *gjd2b* in zebrafish) was identified to be associated with myopia in a recent human GWAS.^6^ Furthermore, a recent study found that knockout of *gjd2a* resulted in hyperopic vision, whereas lack of *gjd2b* led to myopic refraction in zebrafish.^50^ Consistent with these data, retinae from dark-reared fish showed higher *gjd2a* mRNA levels (effect of treatment: *P* = *0.0233*, 2-way ANOVA), especially at 2-week post-treatment (*P* = *0.0397*; Fig. 6K). Surprisingly, there was a remarkable up-regulation of *gjd2b* expression after dark-rearing (effect of treatment: *P* < *0.0001*, 2-way ANOVA; *P* < *0.0001* at 2-week post-treatment; Fig. 6L). These data suggest that altered gap junctions are involved in myopia development induced by dark-rearing in zebrafish. Together these results highlight the complex temporal profile of gene-environment interactions that underlie myopia development in response to adverse visual conditions.

### Dark-rearing induces subcellular retinal changes in myopic zebrafish

In addition to transcriptional analyses using RT-qPCR, we also examined the impact of dark-rearing on post-transcriptional protein expression of EFEMP1, tissue inhibitor of metalloproteinases 2 (TIMP2), and matrix metalloproteinases 2 (MMP2) in the zebrafish retina using immunohistology. As comparing fluorescence levels between immunostained images is questionable, we instead quantified normalized fluorescence intensity (see Materials and Methods section for details of normalization) for analyses of relative protein distribution between inner retinal layers. Although the *efemp1* mRNA level was not significantly changed in zebrafish eyes under dark-rearing (Fig. 6B), our results showed that after 2 days of dark-rearing, inner retinal distribution of EFEMP1 was altered, showing relatively more protein in the amacrine cell layer (ACL; *P* = *0.0019*), but discernibly less in the inner plexiform layer (IPL; *P* = *0.0167*; Fig. 7A–B, 7*i*1, *ii1*, *iii1* and *iv*1). However, after 2 weeks of dark-rearing, we did not find a difference in EFEMP1 distribution (Fig. 7C–D, 7*i*2, *ii*2, *iii*2 and *iv*2).

**Figure 7.**
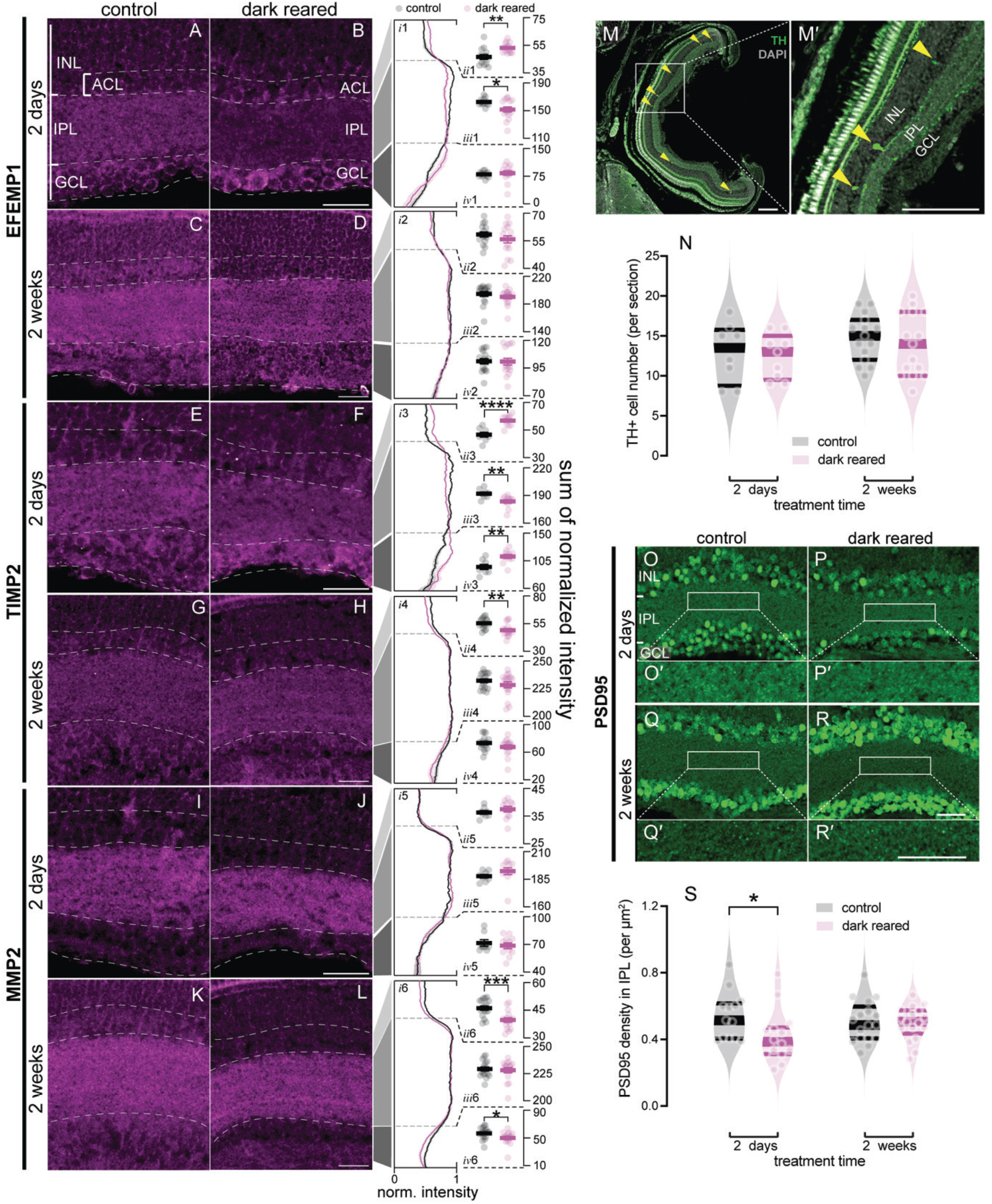
Histological analysis of myopia-associated molecules in the dark-reared zebrafish myopia model. (**A**–**L**) EFEMP1 (**A**–**D**), TIMP2 (**E***–***H)** and MMP2 (**I**–**L**) staining in control (left column) and dark-reared retinae (right column) at 2 days (upper rows) or 2 weeks (lower rows) post-treatment. Normalized expression levels (norm. intensity) for (***i1***– ***i2***) EFEMP1, (***i3****–**i4***) TIMP2 and (***i5****–**i6***) MMP2 across the inner retina in control (black traces) and dark-reared (pink traces) fish at 2 days (upper rows) or 2 weeks (lower rows) post-treatment. Normalized intensity was quantified in the (***ii1****–**ii6***) amacrine cell layer (ACL), (***iii1****–**iii6***) inner plexiform layer (IPL) and (***iv1****–**iv6***) ganglion cell layer (GCL). For EFEMP1, there were 13 control and 15 dark-reared retinae for 2 days post-treatment, and 16 retinae per group for 2 weeks post-treatment. For both TIMP2 and MMP2, there were 8 control and 11 dark-reared retinae for 2 days post-treatment, as well as 17 control and 16 dark-reared retinae for the 2-week dark-rearing timepoint. Data presented as Mean ± SEM. Unpair *t*-test was performed. (**M**) Representative image of dopaminergic amacrine cells (DAC) stained using tyrosine hydroxylase (TH) with a zoomed in region shown in (**M′**) with yellow arrowheads highlight DACs. (**N**) Number of DACs (TH+) was quantified and compared between groups after 2 days and 2 weeks of dark-rearing. For analysis of dopaminergic (or TH+) cell number, there were 8 control and 9 dark-reared retinae for 2 days post-treatment, and 15 control and 15 dark-reared retinae per group for 2 weeks post-treatment. *Two*-way ANOVA with Fisher’s LSD *post-hoc* tests were performed. (**O**–**R′**) PSD95 density in the IPL of control (left column) and dark-reared retinae (right column) with 2 days (upper rows) or 2 weeks (lower rows) post-treatment, with PSD95 also shown at higher magnification (**O′**, **P*′***, **Q′**, and **R′**). (**S**) PSD95 density in the IPL was quantified and compared between groups. For analysis of PSD95 puncta, there were 14 control and 13 dark-reared retinae for 2 days post-treatment, and 17 control and 17 dark-reared retinae for 2 weeks post-treatment. *Two*-way ANOVA with Fisher’s LSD *post-hoc* tests were performed. For all violin plots, dots are data from individual retinae. Thick bars represent medians, and thin lines indicate interquartile ranges. For all statistics, \**P* < *0.05;* \*\**P* < *0.01;* \*\*\**P* < *0.001;* \*\*\*\**P* < *0.0001*.

Activation of MMP2 and increased *TIMP2* mRNA have been reported in myopic eyes.^51–53^ Here, our results showed that after 2 days of dark-rearing, TIMP2 expression was relatively higher in the ACL (*P* < *0.0001*) and ganglion cell layer (GCL; *P* = *0.0041*), but lower in the IPL (*P* = *0.0079*; Fig. 7E–F, 7*i*3, *ii*3, *iii*3 and *iv*3). Conversely, after 2 weeks of treatment, dark-reared fish showed relatively lower TIMP2 expressed in the ACL compared to control fish (*P* = *0.0014*), however, no change was found in the other two layers (Fig. 7G–H, 7*i*4, *ii*4, *iii*4 and *iv*4). For MMP2, there was no discernible distribution changes after 2 days dark-rearing (Fig. 7I–J, 7*i*5, *ii*5, *iii*5 and *iv*5). However, after 2 weeks of dark-rearing, there appeared to be reduced relative expression of MMP2 in the ACL (*P* = *0.0005*) and GCL (*P* = *0.0390*; Fig. 7K–L, 7*i*6, *ii*6, *iii*6 and *iv*6).

A light-driven decrease in dopamine (or increased dopamine turnover) levels has been linked to myopia development.^54, 55^ As dopaminergic amacrine cells (DAC) are the main source of retinal dopamine,^56^ DAC in zebrafish retinae were labelled by tyrosine hydroxylase (TH) and cell density was compared (Fig. 7M–M′, yellow arrowheads). There was no change in retinal DAC (or TH+ cell) density in dark-reared zebrafish compared with controls (Fig. 7N)

It has been shown that EGR1 inhibits *post-synaptic density 95* (*PSD95*) transcription in response to activation of the *N*-Methyl-d-aspartate receptor (NMDAR) in the mouse hippocampus.^57^ Given the reduction of *egr1* (Fig. 6A) in dark-reared zebrafish, we quantified retinal PSD95 expression, expecting PSD95 density to be increased in the IPL after dark-rearing. However, we found that after 2 days of dark-rearing, zebrafish retinae had lower PSD95 density than control fish (*P* = *0.0191*; Fig. 7O–P′ and 7S), and after 2 weeks of dark-rearing there was no difference between groups (Fig. 7Q–R′ and 7S). Therefore, our results suggest that EGR1 regulates retinal PSD95 differently to the brain.

## Discussion

Here, we showed that 2-week dark-rearing of 4-week-old zebrafish is a reliable means of myopia induction (axial and perhaps equatorial, Fig. 1C and 3B–C). This phenotype is recoverable if dark-rearing does not exceed 2 weeks (Fig. 3B–C). The establishment of a robust environmental zebrafish myopia model may be a boon for high-throughput studies of myopia mechanisms and interventions. Importantly, with a range of well-established gene editing tools available for zebrafish, tools not available for other species except mice, this model is an excellent *in vivo* platform for studying gene-environment interactions in myopia development.

Approaches to environmental myopia induction including diffusor or negative lens wearing are not feasible in zebrafish. Besides dark-rearing, rearing in restricted wavelengths light may also be used for myopia induction in zebrafish. In other animals like chicks, guinea pigs and squid, long-wavelength (641, 530 and 557 nm, respectively) rearing causes myopia, whereas short-wavelength (477, 430 and 447 nm, respectively) rearing inhibits eye elongation, leading to hyperopia.^16, 58, 59^ Intriguingly, the same wavelength bands produce the opposite refractive phenotypes in tree shrews.^60, 61^ Consistent with the tree shrew, long-wavelength exposure (above 570 nm) imposes hyperopia in rhesus monkeys.^62^ Moreover, a recent study found that restricted light spectra impacts zebrafish ocular growth, with rearing under cyan (483 nm) or red light (633 nm) both promoting hyperopia in zebrafish.^63^ These results point to a species-dependent response to restricted-wavelength rearing for ocular development, which is likely to reflect evolutionary adaptations of visual systems to their living environments. Finally, whilst as yet not attempted, modification of spatial or temporal characteristics of visual input may also induce myopia in zebrafish. Examples include myopia development in monkeys reared in a restricted visual space,^64^ and in guinea pigs exposed to 0.5 Hz flicker light.^65^ Manipulating brightness, wavelength, as well as spatial and temporal features of the visual environment in zebrafish can help identify the relative importance of these factors in myopia development.

Our ERG results indicate changes in both photoreceptor and bipolar cell function in zebrafish even after only 2 days of dark-rearing (Fig. 5A–E). Attenuated ERG responses have been noted in myopic eyes in human clinical studies.^66^ During myopia development, excessive eye elongation can lead to retinal current sources being farther from corneal electrodes, and stretching and damage to the retina, might all contribute to smaller ERG responses. Here we show that functional deficits preceded presence of myopia-associated structural changes, and thus may indicate that altered retinal signaling is an early event in myopia development. Moreover, faster *b*-waves after 2 weeks of dark-rearing (Fig. 5F–J) points to changes to inner retinal processing and argues against passive structural changes, which should only attenuate amplitude. It is noteworthy that dark-rearing is a relatively extreme disruption to the visual environment, and may thus have more impact on retinal function. Indeed, ERG responses are not altered in mice with lens-induced high myopia.^67^

Using OMR, we demonstrated visual behavioral deficits in 2-week dark-reared zebrafish (Fig. 4A–B). Our results correspond with reported human data, which showed reduced contrast sensitivity at high spatial frequencies in high myopia.^68^ OMR deficits in dark-reared zebrafish can result from impaired retinal function or refractive error. Future studies might separate these potential causes by considering behavioral function earlier in this model.

Our qPCR and histological characterizations provided a molecular basis for dark-rearing-induced altered retinal function. Down-regulated *vefgaa* and *vegfab* expression in 2-day dark-reared zebrafish eyes (Fig. 6E–F) corresponds with a previous report that retinal pigmented epithelial-specific *VEGFA* knockout in mice disrupts cone function.^69^ Down-regulation was transient, suggesting involvement of *Vegfas* in the early functional changes in this myopia model. Down-regulation of *rbp3* (Fig. 6J), a key gene in the visual cycle and phototransduction,^70^ might contribute to dark-rearing-induced reduced ERG responses. In addition, we found reduced density of PSD95, a crucial excitatory post-synaptic scaffolding protein, in the IPL of 2-day dark-reared zebrafish (Fig. 7O–S). There was no difference in PSD95 density after 2 weeks of dark-rearing, suggesting existence of other contributors to the functional deficits at this later time point.

In our gene expression analyses, *egr1* down-regulation in dark-reared zebrafish (Fig. 6A) is consistent with findings from other environmental myopia models.^35–37^ We anticipated that *egr1* down-regulation would lead to up-regulation of PSD95, as has been reported in other tissues.^57^ However, we observed the converse, a PSD95 down-regulation in zebrafish that requires further investigation.

Expression changes of gap junction genes, *gjd2a* and *gjd2b*, suggest that they play roles in myopia development in zebrafish (Fig. 6K–L). Consistent with previous results showing that *gjd2a* knockout induces hyperopia in zebrafish,^50^ *gjd2a* up-regulation was observed in our dark-reared model, which suggests involvement of *gjd2a* in the directional control of ocular refraction. Given that *gjd2b*-knockout zebrafish developed myopia,^50^ one would have expected a *gjd2b* down-regulation in our dark-reared zebrafish. Indeed, decreased Connexin36 (the mammalian orthologue of *gjd2*) content in the inner retina of form-deprived myopic guinea pigs has been reported.^71^ However, this was not the case in our study; we showed that *gjd2b* was unexpectedly up-regulated in 2-week dark-reared zebrafish. We hypothesized that *gjd2b* up-regulation in zebrafish may be triggered by an intrinsic feedback loop modulating *gjd2b* mRNA levels for non-environmental growth regulation. Intrinsic regulation of zebrafish ocular refraction has been previously observed; dark-reared zebrafish developed myopia within the first month, but longer dark-rearing produced evidence of recovery.^14^ Mice lacking *connexin36* showed impaired synaptic transmission in the rod pathway, with reduced scotopic *b*-wave amplitude.^72^ However, increased *gjd2s* expression in dark-reared zebrafish, which should increase connexins, did not rescue their attenuated *b*-waves, suggesting that retinal functional defects driven by dark-rearing involve a number of different pathways, *rbp3* for example. Therefore, up-regulated *gjd2s* alone might not be sufficient to compensate for retinal impairment. Mechanistic insights into gap junction-associated myopia development in a range of models will be an interesting subject for further study.

Our qPCR data did not indicate involvement of *efemp1*, *tgfb1s*, *igf1*, *fgf2* and *wnt2b* in dark-rearing-induced myopia in zebrafish. It is worth noting that firstly we analyzed mRNA isolated from the whole zebrafish eye, which may account for differences between ours and previous results. Second, the myopia-risk genes tested here were identified by human GWAS or from other animal models with form-deprivation or lens-induced myopia. Given differences due to tissue sampling, species and modes of myopia induction, differences in gene expression profile may not be surprising.

We found altered distribution of EFEMP1, TIMP2 and MMP2 in dark-reared zebrafish retinae (Fig. 7). Changes in relative distribution were layer and time dependent. After 2 days of dark-rearing, relative expressions of EFEMP1 and TIMP2 were decreased in the IPL but increased in the ACL and/or GCL. By 2 weeks post-dark-rearing TIMP2 was decreased in the ACL, and MMP2 was decreased in both ACL and GCL. Whilst these results require further investigation, they emphasize that in addition to changes to mRNA expression, the distribution of relevant molecules is important to consider when investigate their function.

We also attempted to determine whether dark-rearing impacts the dopamine pathway, a long-standing pathway associated with myopia.^73^ However, our result showed that DAC number were unchanged for dark-reared zebrafish. It is worth noting that the amount of dopamine released or its turnover in the eye can be impacted without changing the DAC number; immediate changes to dopamine release and turnover in response to 1 hour/1hour light/dark pulse have been shown in the mouse retina.^74^

In summary, two weeks of dark-rearing produces a robust but recoverable myopia in zebrafish, with a phenotype that manifests at many levels, including impaired visual sensitivity, altered retinal function, and changes in expression and distribution of many myopia-associated molecules, such as EGR1, VEGFAs, RBP3, GJD2s, EFEMP1, TIMP2, MMP2 and PSD95. With the advantages afforded by large-scale genetic editing tools in this species, this environmental zebrafish myopia model will facilitate high-throughput *in vivo* investigation of gene-environment interactions currently not available to other myopia models. Together with the multi-level assessment illustrated in the present study, we believe that this myopia model will be a valuable platform for in-depth examination of the growing number of myopia-risk genes.

## Acknowledgments

We thank Yoshi Itoh for his kind gift of a sheet anti-TIMP2 antibody. We thank the Biological Optical Microscopy Platform (BOMP) at the University of Melbourne for providing instrument for confocal imaging. We acknowledge the staff of the Danio rerio University of Melbourne and Walter and Eliza Hall of Medical Institute fish facilities for animal husbandry and support.

JX was supported by the Melbourne Research Scholarship for his PhD study and the Dr Albert Shimmins Postgraduate Writing-Up Award for manuscript preparation. Funding was provided through the University of Melbourne research grant support scheme.

## Commercial relationship

**J. Xie**, None; **P.T. Goodbourn**, None; **B.V. Bui**, None; **P.R. Jusuf**, None.

